# Explicit and implicit motor simulations are impaired in individuals with aphantasia

**DOI:** 10.1101/2022.12.15.520602

**Authors:** W. Dupont, C. Papaxanthis, C. Madden-Lombardi, F. Lebon

**Affiliations:** INSERM UMR1093-CAPS, Université Bourgogne Franche-Comté, UFR des Sciences du Sport, F-21000, Dijon; Centre National de la Recherche Scientifique (CNRS), France; Institut Universitaire de France (IUF)

**Keywords:** Motor imagery, Action observation, Aphantasia, Mental representation

## Abstract

Individuals with aphantasia report having difficulties or an inability to generate visual images of objects or events. So far, there is no evidence showing that this condition also impacts the motor system and the generation of motor simulations. We probed the neurophysiological marker of aphantasia during explicit and implicit forms of motor simulation, i.e., motor imagery and action observation, respectively. We tested a group of individuals without any reported imagery deficits (phantasics) as well as a group of individuals self-reporting the inability to form mental images (aphantasics). We instructed the participants to explicitly imagine a maximal pinch movement in the visual and kinesthetic modalities and to observe a video showing a pinch movement. By means of transcranial magnetic stimulation, we triggered motor-evoked potentials (MEP) in the target right index finger. As expected, MEP amplitude, a marker of corticospinal excitability, increased for phantasics during kinaesthetic motor imagery and action observation relative to rest but not during visual motor imagery. Interestingly, MEP amplitude did not increase in either form of motor simulation for the group of aphantasics. This result provides neurophysiological evidence that individuals living with aphantasia have a real deficit in activating the motor system during motor simulations.

## Introduction

The generation of mental simulations is a fundamental characteristic of human existence, allowing us to retrieve and/or predict the sensorimotor consequences of an action (motor imagery) or to understand the actions of others (action observation). Motor imagery and action observation, also identified as explicit and implicit motor simulations, respectively (Jeannerod, 2001), activate the sensorimotor system, although no movement is produced (Caspers et al., 2010; Courson and Tremblay, 2020; Hardwick et al., 2018; Hétu et al., 2013; Iacoboni et al., 2005; Ruffino et al., 2017; Sharma et al., 2008). Such mental processes are relevant interventions to improve motor learning and to promote motor rehabilitation (Malouin et al., 2013; Rannaud Monany et al., 2022; Ruffino et al., 2021, 2017).

Although the generation and use of mental simulation seems a natural process, a small portion of the population, called aphantasics, report being unable or struggling to create mental images of an event, although they do not have associated disorders (Dawes et al., 2020; Milton et al., 2021; Zeman et al., 2020, 2015). Behaviorally, this deficit is mainly evaluated by subjective reports of visual imagery vividness (Dawes et al., 2020; Milton et al., 2021; Zeman et al., 2020, 2016, 2015, 2010). Kay and colleagues recently used physiological methodologies, such as binocular rivalry and eye tracking, to show that aphantasics have difficulty visualizing events (Kay et al., 2022 and Keogh and Pearson, 2018). However, the available psychological and physiological evidence at present remains insufficient to determine whether aphantasics are actually unable to generate mental simulations or whether such difficulty is a matter of strategy or metacognition. In addition, it remains unclear whether this deficit that is typically measured in visual imagery would also affect motor imagery.

The present paper aims to shed new light on aphantasics’ condition by exploring their ability to activate the motor system while engaging in an explicit form of motor simulation, i.e., motor imagery, using neurological measures that do not rely on self-report. It is, however, possible that any observed psychophysical or neurophysiological deficits could be related to individual differences in strategies or motivation (“I don’t think I can imagine, so I don’t try”). Therefore, we also tested the modulation of the motor system during action observation, which engages an implicit form of motor simulation, unrelated to the participant’s efforts or strategies. For each of these simulation types, we measured corticospinal excitability by means of Transcranial Magnetic Stimulation (TMS) as well as subjective reports of imagery vividness. There is strong evidence in the literature that neurophysiological manifestations of motor simulations in individuals with normal imagery ability (i.e., phantasics) include the increase of corticospinal excitability in TMS studies during both motor imagery (Facchini et al., 2002; Fadiga et al., 1998; Grosprêtre et al., 2016; Lebon et al., 2012; Rossini et al., 1999; Ruffino et al., 2017) and action observation (Aziz-Zadeh et al., 2002; Borroni et al., 2005; Brighina et al., 2000; Clark et al., 2004; Fadiga et al., 2005, 1995; Gangitano et al., 2001; Strafella and Paus, 2000).

Our main hypothesis is straightforward: if aphantasics are not able to create motor simulations (self-reports), we would not observe an increase of corticospinal excitability during motor imagery and action observation that is typically measured in phantasics. The absence of motor cortex activation during motor simulations would be a relevant marker of aphantasia.

## Results

### Explicit motor simulation (motor imagery)

Aphantasics (n=15) reported having difficulties or even the inability to explicitly create mental images of common actions, in comparison to phantasics (n=15). Average scores on the Vividness of Movement Imagery Questionnaire-2 (Roberts et al., 2008) and at the Spontaneous Use of Imagery Scale (Ceschi and Pictet, 2018) approached the boundary of no imagery for aphantasics (see Figure 1 for statistics and main results and Table 1 in supplementary section for details). These subjective reports are in line with the literature focusing on the creation of mental visual images (Dawes et al., 2020; Keogh and Pearson, 2018; Zeman et al., 2015).

**Figure 1:**
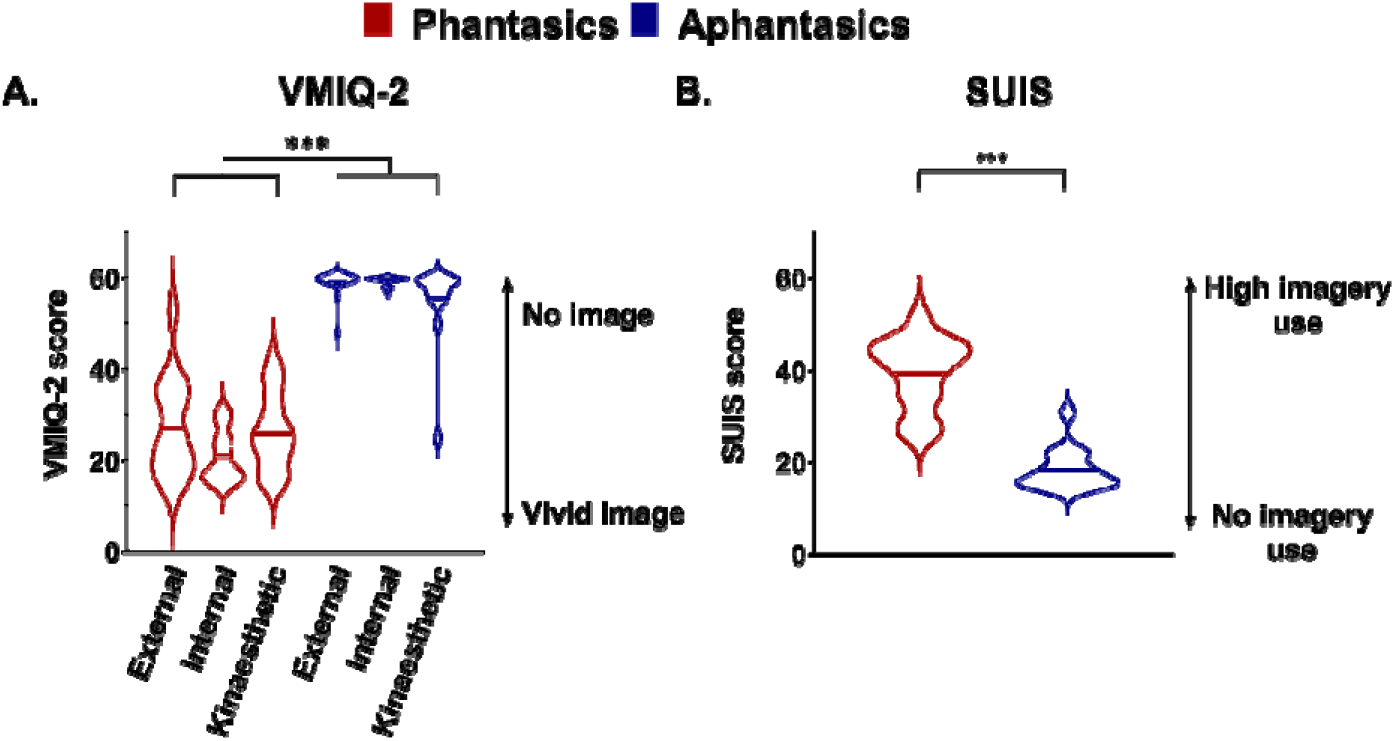
Violin plots represent scores on **A**) the Vividness of Movement Imagery Questionnaire-2 (VMIQ-2) and **B**) the Spontaneous Use of Imagery Scale (SUIS) questionnaire for phantasics and aphantasics. Thick horizontal lines mark the mean, and circles represent individual data. For VMIQ-2, lowest (12) and highest (60) scores represent the highest and lowest vividness, respectively. The participants reported the vividness of imagined movements for three modalities (external visual, internal visual and kineasthetic). The scale is inverted for the SUIS, where the lowest (12) and highest (60) scores represent lowest and highest use of mental imagery in everyday situations, respectively. *** = p<0.001.

These subjective reports were supplemented by neurophysiological data. We delivered single-pulse TMS over the finger/hand muscle area of the left primary motor cortex while participants were at rest and while they imagined a maximal pinch movement in the visual or kinaesthetic modality (what the movement looks like, or how it feels, respectively). Corticospinal excitability was probed in the first dorsal interosseus muscle via the peak-to-peak amplitude of the motor-evoked potential (MEP). As expected, corticospinal excitability increased during kinaesthetic but not visual motor imagery in phantasics (Stinear et al., 2006), whereas it was not modulated in either imagery modality for aphantasics (Figure 2). More specifically, MEP amplitude increased during kinaesthetic imagery in comparison to rest for phantasics (24.15 ±25.58%; t=3.65, p=0.002), but not for aphantasics (−2.01 ±46.88%; t=-0.166, p=0.870). The percentage of MEP amplitude increase differed between the 2 groups (one-sided t-test: t(_28_)=1.897, p=0.034; Cohen’s d=0.72). During visual imagery, neither group increased MEP amplitude in comparison to rest (−5.29 ±27.88%, t=-0.735, p=0.474, and -1.33 ±31.75%, t=-0.162, p=0.873 for phantasics and aphantasics, respectively). The percentage of MEP amplitude increase was not statistically different between the 2 groups (one-sided t-test: t(_28_)=-0.363, p=0.719; Cohen’s d=0.14).

**Figure 2:**
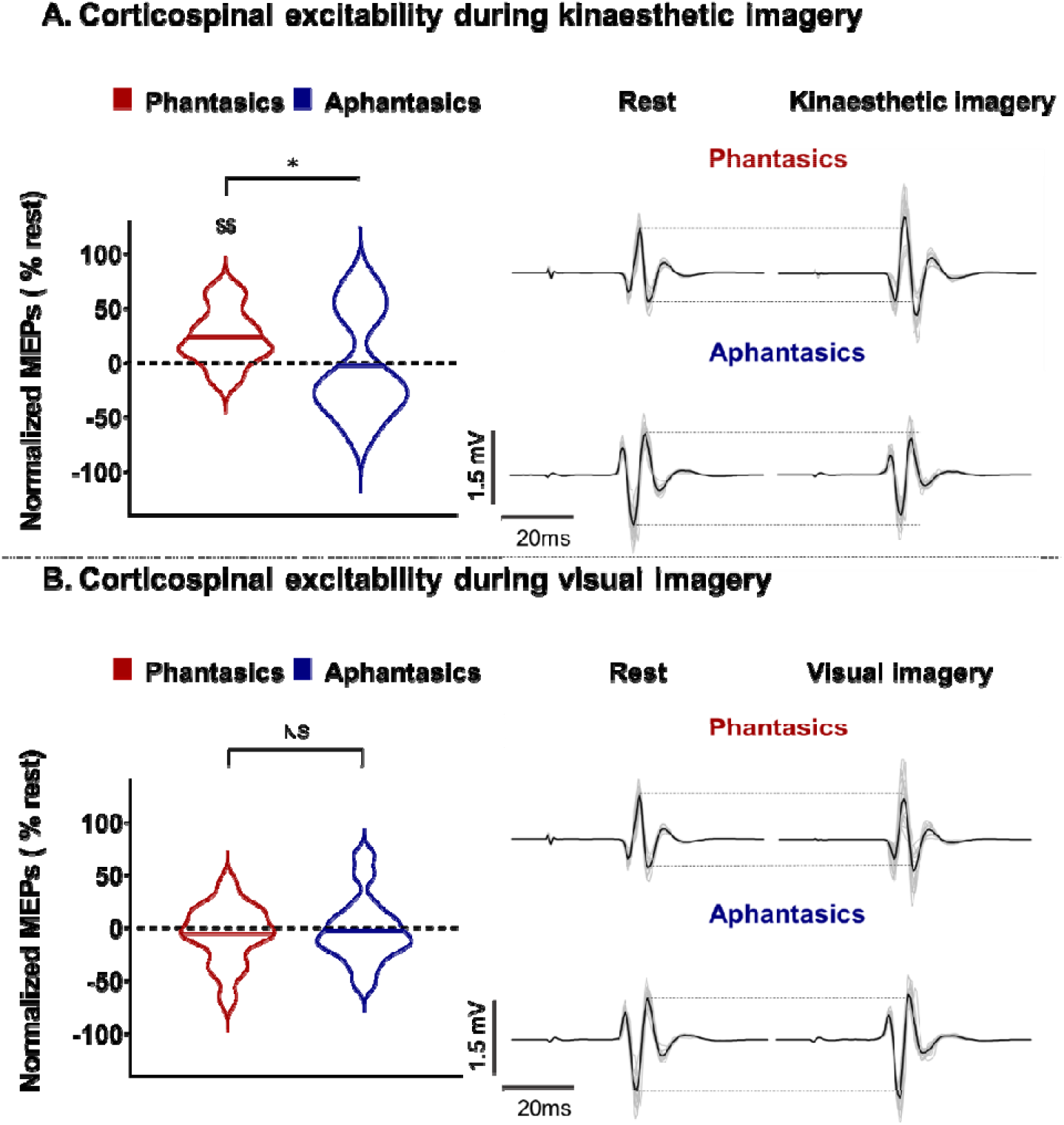
Corticospinal excitability during kinaesthetic and visual imagery for phantasics and aphantasics. Violin plots on the left side represent normalized MEPs during kinaesthetic (**A**) and visual imagery (**B**). Thick horizontal lines mark the mean. Circles represent individual data. The right side of the panel illustrates raw MEPs of a typical subject in grey lines, and the black line is the average MEP of the condition for this participant. *=p<0.05 indicates a significant difference between the two groups and $$=p<0.01 indicates a significant difference from zero (rest).

### Implicit motor simulation (action observation)

While the previous kinaesthetic imagery measure provides strong evidence that aphantasics do not explicitly generate motor images when prompted, it remains possible that they are capable of simulating, but avoid doing so, perhaps due to their impressions of difficulty or failure. Therefore, we also measured corticospinal excitability during an implicit form of motor simulation that is less influenced by participants’ efforts and strategies. Participants observed a pinch movement of the right hand on a computer screen (see method section for details on TMS timings). As expected, MEP amplitude increased during action observation in comparison to rest for phantasics (13.89 ±24.28%, t=2.215, p=0.043). Interestingly, MEP amplitude decreased for aphantasics (−14.95 ±23.46%, t=-2.469, p=0.027). The percentage of MEP amplitude increase was statistically different between the 2 groups (one-sided t-test: t(_28_)=3.309, p=0.002; Cohen’s d=1.25; Figure 3).

**Figure 3:**
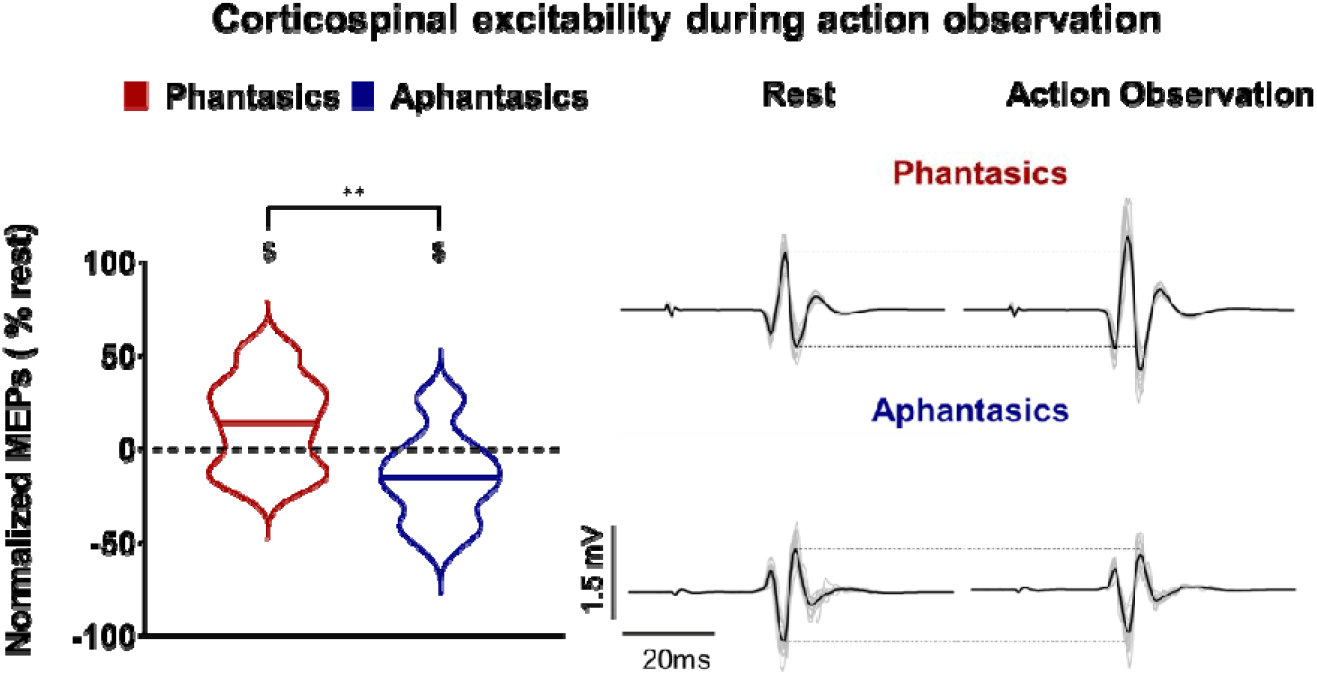
Corticospinal excitability during action observation for phantasics and aphantasics. Violin plots on the left side represent normalized MEPs during action observation. Thick horizontal lines mark the mean. Circles represent individual data. The right side of the panel illustrates raw MEPs of a typical subject in grey lines, and the black line is the average MEP of the condition for this participant. **=p<0.005 indicates a significant difference between the two groups and $=p<0.05 indicates a significant difference from zero (rest).

## Discussion

The present study provides several notable findings related to motor simulation and aphantasia. First, the increase in corticospinal excitability during explicit motor imagery for phantasics but not aphantasics demonstrates that only those able to generate mental simulations of hand actions showed motor cortex engagement. Second, this same effect during action observation, an implicit form of motor simulation, demonstrates once again that aphantasia entails a real impairment outside the control of individuals (strategies, volition, metacognition).

Concerning the first point, the self-reported estimates of vividness (VMIQ-2 and SUIS) suggest that participants living with aphantasia are less able (or unable) to simulate movements, corroborating previous research on mental imagery (Dawes et al., 2020; Keogh and Pearson, 2018; Zeman et al., 2015). However, it remains unclear whether such reports reflect a real neurophysiological impairment, or merely a metacognitive deficit. Thus, we used the modulation of corticospinal excitability as a marker of the generation of motor simulation. We confirmed the increase of corticospinal excitability during kinaesthetic imagery in phantasics (e.g., Facchini et al., 2002; Fadiga et al., 1998; Grosprêtre et al., 2016; Kasai et al., 1997; Rossini et al., 1999; Ruffino et al., 2017) and the absence of such an increase in the motor system during visual imagery (Stinear et al., 2006). More interestingly, we found that aphantasics did not exhibit any increase of corticospinal excitability during imagery in either modality. These psychological and physiological findings support the view that aphantasia entails a real impairment in the generation of motor images, rather than a metacognition failure (Kay et al., 2022; Keogh and Pearson, 2021, 2018). To note, our sample of aphantasics was self-selected (volunteers who identified themselves as unable to generate imagery), and our phantasic sample was not pre-screened for imagery ability. Therefore, our groups may not reflect the maximal difference between phantasic and aphantasic individuals. Specifically, some of our aphantasics may not be fully impaired in their simulation ability, but rather possess only a limited ability to simulate (Zeman et al., 2020). Our data also lends support to the idea that certain aphantasics can present complete impairments in one modality (e.g., visual) while exhibiting at least limited abilities to simulate in another modality (e.g., kinaesthetic) (Dawes et al., 2020).

Concerning the second point, the current study is, to our knowledge, the first to illustrate that aphantasia also encompasses an inability to engage in implicit motor simulations. This result extends recent investigations demonstrating the deterioration of implicit simulations during mental rotation (Pounder et al., 2022; Zeman et al., 2010; Zhao et al., 2022; but Milton et al., 2021) and involuntary simulations during night-time dreams (Dawes et al., 2020; Zeman et al., 2015, 2010; see review: Whiteley, 2020). As already described for motor imagery, we observed that, relative to rest, action observation increased corticospinal excitability in phantasics but not aphantasics. Indeed, in the case of aphantasia, the reported absence of a motor simulation is coupled with a real deficit, manifest in the lack of an increase in corticospinal excitability during action observation.

To conclude, our results provide novel neurophysiological evidence that aphantasia is marked by a measurable lack of activation in the motor system, even when motor simulation should be engaged implicitly rather than explicitly. The modulation of MEP amplitude during explicit and implicit motor simulation may be a relevant tool to characterize aphantasia in the motor domain.

## Material and methods

### Participants

Thirty-four right-handed aphantasic (n=17) and phantasic (n=17) participants were included in the study. Four participants with extreme values were excluded from the analysis (see Data and statistical analysis for details). We performed the statistical analyses with 15 aphantasics (9 women, mean age: 20; range: 18-26) and 15 phantasics (6 women, mean age: 23; range: 19-26). We ensured right laterality with the Edinburgh inventory (Oldfield, 1971). All participants were French native speaker and completed the questionnaire by Lefaucheur et al., 2011 prior to participation to determine whether they were eligible for TMS. All procedures (excluding pre-registration) were approved by an ethics committee (CPP SOOM III, ClinicalTrials.gov Identifier: NCT03334526) and were in accordance with the Declaration of Helsinki.

### Procedure and stimuli

Participants came to the laboratory for two experimental sessions. First, the behavioral session, included subjective assessments of imagery. Then, the neurophysiological session, consisted of measures of corticospinal excitability by means of single-pulse TMS delivered at rest, during visual and kinaesthetic motor imagery and during action observation.

### Behavioral session

All participants completed the Vividness of Movement Imagery Questionnaire-2 (VMIQ-2) (Roberts et al., 2008) and the Spontaneous Use of Imagery Scale (SUIS) (Ceschi and Pictet, 2018). In the VMIQ-2, participants imagined multiple actions in three modalities (External Visual Imagery, Internal Visual Imagery, Kinaesthetic imagery), and then rated how vivid their imagery was for each movement on a scale of 1-5 (with 1 = “Perfectly clear and vivid as normal vision” and 5= “No image at all, you only think about the movement”). The SUIS evaluated the general tendency of individuals to use visual mental imagery in everyday situations. Each item is rated on a 5-point scale (from 1 = “never appropriate”; to 5 = “always completely appropriate”) according to the item described and the individual’s functioning.

### Neurophysiological session

In this session, we used TMS to probe corticospinal excitability at rest, during visual and kinaesthetic motor imagery and during AO. Participants sat in front of a 19-inch LCD monitor. First, sixteen TMS pulses were delivered at rest (fixation cross), which served as baseline.

In each motor imagery block, participants were instructed to imagine sixteen maximum voluntary contractions of pinch movements either in a 1^st^-person visual or kinaesthetic modality, corresponding to the experimental conditions Visual Imagery and Kinaesthetic Imagery, respectively. The following instructions were provided: “try to imagine yourself performing the pinch movement, by visualizing the movement just as if you were watching your fingers move (for visual imagery) or feeling the fingers sensations as if you were doing the movement (for kinaesthetic imagery)”. TMS pulses were delivered 2000 ms after the cue to imagine (“O” on the screen). The inter-trial interval was 7000ms. In the action observation block, participants were instructed to observe a video of a pinch movement. For each of the sixteen trials, a TMS pulse was delivered 1000 ms after the touch between the index and the thumb fingers. The blocks were counterbalanced between participants.

Surface electromyography (EMG) was recorded using 10-mm-diameter surface electrodes (Contrôle Graphique Médical, Brice Comte-Robert, France) placed over the right First Dorsal Interosseous (FDI) muscle. Before positioning the electrodes, the skin was shaved and cleaned to reduce EMG signal noise (<20μV). EMG was amplified with a bandpass filter (10-1000 Hz) and digitized at 2000Hz (AcqKnowledge; Biopac Systems, Inc., Goleta, CA). We calculated the root mean square EMG signal (EMGrms) for further analysis.

Using a figure-eight coil (70 mm diameter) connected to a Magstim 200 stimulator (Magstim Company Ltd, Whitland), single-pulse TMS was delivered over the motor area of the right FDI muscle. The coil rested tangentially to the scalp with the handle pointing backward and laterally at a 45° angle from the midline. Using a neuronavigation system (BrainSight, Rogue Research Inc.) with a probabilistic approach (Sparing et al., 2008), the muscle hotspot was identified as the position where stimulation evoked the highest and most consistent MEP amplitude for the FDI muscle. This position was determined via a regular grid of 4 by 4 coil positions with a spacing of 1 cm (centered above the FDI cortical area x=-37, y=-19, z=63; Bungert et al., 2017; Sondergaard et al., 2021). During the experiment, the intensity of TMS pulses was set at 130% of the resting motor threshold, which is the minimal TMS intensity required to evoke MEPs of 50µV peak-to-peak amplitude in the right FDI muscle for 5 out of 10 trials (Rossini et al., 2015).

### Data and statistical analysis

Matlab (The MathWorks, Natick, Massachusetts, USA) was employed to extract EMG and we extracted the peak-to-peak MEP amplitude. Before statistical analysis, we discarded MEPs outside the range of +/-2 SDs from individual means for each condition (3.75% of all data). We normalized the average MEP amplitude for each condition to rest. Four participants (two phantasics and two aphantasics) were removed from the final analysis due to extreme values (outside the range of 2 SDs). Using, Shapiro-Wilk and Mauchly tests, we checked the normality and sphericity of the data. One-sided t-tests were used to compare MEPs between each group during kinaesthetic and visual imagery, and action observation. Moreover, one-sample t-tests were used to compare condition MEPs to zero (rest). Finally, to ensure that MEPs were not biased by muscle activation preceding stimulation, we used a Friedmann ANOVA to compare the EMGrms before the stimulation artifact in all our conditions for each group (See Supplementary section). Statistics and data analyses were performed using the Statistica software (Stat Soft, France). The data are presented as mean values (±SD) and the alpha value was set at 0.05.

## Supporting information

Supplemntary results

## Data availability statement

All data from this study are available at:

https://osf.io/4apxw/?view_only=f3dd901d53424462beaa8926cbda6dcc

