## Supplementary material for "Explicit and implicit motor simulations are impaired in individuals with aphantasia": Supplemntary results

**Supplementary results**

Table 1: Average scores on the Vividness of Movement Imagery Questionnaire-2 (VMIQ-2, Roberts et al., 2008) and the Spontaneous Use of Imagery Scale (SUIS, Ceschi and Pictet, 2018) for aphantasic and phantasic participants. The worst and best score for each modality at the VMIQ-2 is 60 and 12, respectively. The worst and best score for the SUIS questionnaire is 12 and 60, respectively. A repeated-measures ANOVA on VMIQ-2 scores revealed a main effect of Group (F_1,28_=324.04, p<0.001, ηp²=0.920), with greater imagery ability for Phantasic (24.91 ±8.86; Cohen’s d=4.60) than Aphantasic participants (57.95 ±5.64). We observed an interaction between Group and Perspective (F_2,56_=4.136, p=0.021, ηp²=0.128), demonstrating a marginally significant difference between the internal (21.06 ±5.59) and external (26.26 ±8.51, p=0.085, Cohen's d=0.75) vividness of visual motor imagery in phantasics, and no difference between perspectives in aphantasics. Moreover, an independent T-test on SUIS scores yielded a greater utilization of imagery in everyday life for Phantasic individuals (39.60 ±7.83) than that for Aphantasic participants (18.53 ±4.47; t(_28_)=9.04, p<0.001, Cohen’s d=3.42).

| Questionnaires | VMIQ-2 | | | SUIS |
| --- | --- | --- | --- | --- |
|  | External visual | Internal visual | Kinaesthetic |  |
| Phantasics | 27.40  ±10.89 | 21.06  ±5.59 | 26.26  ±8.51 | 39.60  ±7.83 |
| Aphantasics | 58.93  ±3.19 | 59.40  ±1.05 | 55.53  ±8.90 | 18.53  ±4.47 |

Table 2: Average scores on mental arithmetic and Raven's matrix for aphantasic and phantasic participants. Using Independent T-tests on mental arithmetic and raven’s matrix scores, no difference was observed between Phantasics and Aphantasics participants on either task.

| Tasks | Arithmetic | Raven’s matrix |
| --- | --- | --- |
| Phantasics | 75.70 ±17.66 | 83.32 ±14.45 |
| Aphantasics | 64.28 ±27.40 | 79.04 ±16.74 |
| T-tests | 0.18 (t=1.35) | 0.45 (t=0.75) |

**EMGrms:**

The absence of muscular pre-activity during action reading and motor imagery was confirmed by a Friedman Anova revealing no significant difference in EMGrms before the TMS artefact between rest, Visual Imagery, Kinaesthetic Imagery and Action Observation conditions for Aphantasic (p=0.430; r=-0.005) and Phantasic individuals (p=0.171, uncorrected; r=0.047).

Table 3: EMGrms activity (mean ±SD) in µV recorded for the first dorsal interosseous before the TMS artifact for each condition (window of 100ms prior the artifact).

|  | Rest | Action observation | Visual Imagery | Kinaesthetic Imagery |
| --- | --- | --- | --- | --- |
| Phantasics | 1.687  ±1.867 | 1.029  ±0.266 | 0.963  ±0.127 | 1.678  ±1.588 |
| Aphantasics | 1.746  ±1.533 | 1.166  ±0.361 | 1.584  ±0.739 | 1.632  ±0.992 |
